# Differential regulation of p62-ubiquitin conjugates in neurons versus astrocytes during cellular stress

**DOI:** 10.1101/2025.11.17.688722

**Authors:** David K. Sidibe, Erin M. Smith, Maeve L. Spivey, Maria C. Vogel, Sandra Maday

## Abstract

Sequestosome 1/p62 (hereafter referred to as p62) is a multifunctional protein that orchestrates various cellular stress response pathways including autophagy, proteasome-mediated degradation, antioxidant defense, nutrient sensing, and inflammatory signaling. Mutations in distinct functional domains of p62 are linked with the neurodegenerative disease amyotrophic lateral sclerosis (ALS), underscoring its importance in neural cells. Neurons and astrocytes perform distinct roles in brain physiology and thus encounter a unique landscape of cellular stress. However, how p62 is regulated in these cell types in response to various stress modalities remains largely unexplored. Several functions for p62 depend on engagement with ubiquitinated substrates. Thus, we investigated how the regulation of p62-ubiquitin conjugates differs between neurons and astrocytes exposed to two stress modalities: lysosomal membrane damage and metabolic stress. Lysosomal damage triggered ubiquitin-dependent assembly of p62 puncta in both neurons and astrocytes. In contrast, nutrient deprivation elicited different responses between neurons and astrocytes. Neurons formed p62-ubiquitin structures more prominently and displayed a greater dependence on ubiquitin for p62 clustering. Together, these findings reveal cell-type-specific and stress-specific regulation of p62-ubiquitin conjugates, indicating that neurons and astrocytes can deploy distinct quality control strategies.

## INTRODUCTION

Sequestosome 1 or p62 (hereafter referred to as p62) is a multifunctional protein that is extensively studied for its role as a receptor for selective autophagy [1–3]. P62 binds ubiquitinated substrates via its ubiquitin-associated domain (e.g., UBA domain) and can bridge cargoes with the autophagy machinery through its LC3-interacting region (e.g., LIR) [1]. P62 also has the capacity to self-oligomerize and assemble into phase-separated condensates, which may facilitate many of the cellular functions of p62 [4–8]. With these properties, p62 promotes autophagic clearance of protein aggregates and organelles to maintain cellular homeostasis [1–3, 9–12].

Beyond canonical roles in autophagy, p62 also plays important roles in the ubiquitin-proteasome pathway [13, 14], anti-oxidant signaling [15–17], pro-inflammatory signaling [18–20], nutrient sensing [21–23], and apoptosis [1]. Thus, p62 integrates protein and organelle quality control and stress response pathways. Many of these functions rely on p62 engaging ubiquitinated substrates to be routed for degradation via autophagosomes [1–3, 24–26]. The critical importance of p62 is evidenced by mutations in p62 that are associated with diseases that manifest differently across tissues, including the neurodegenerative disorder amyotrophic lateral sclerosis (ALS) [27–29] and the degenerative bone disease, Paget’s disease [30, 31]. Despite extensive study of p62 in immortalized cell lines, the mechanistic diversity of p62 pathway engagement in key cell types of the brain, such as neurons and astrocytes, remains poorly understood.

Neurons and astrocytes perform distinct functions in the brain. Neurons are specialized for rapid synaptic transmission and astrocytes regulate synaptic function, provide metabolic support and serve key roles in buffering oxidative insults and coordinating immune responses. Through these distinct roles in brain physiology, each cell type encounters a unique landscape of cellular stress, which may necessitate different adaptations within quality control and stress response pathways. Indeed, we have shown that metabolic stress and proteotoxic stress induced by proteasomal inhibition elicits differential responses by the autophagy-lysosomal pathway in neurons versus astrocytes [32, 33]. Moreover, Rhoads et al. demonstrated that neurons and astrocytes display distinct organelle signatures, encompassing organelle density, dynamics, and inter-organelle contacts, that shift in a stress-dependent manner [34]. However, how p62 may integrate these divergent stress responses in a cell-type-specific manner remains an open question.

Here, we employ a neuron-astrocyte coculture platform to define stress- and cell-type-specific regulation of p62 and its engagement with ubiquitinated substrates. We focus on two stress paradigms that induce quality control pathways previously linked with p62 engagement: lysosomal membrane damage and nutrient deprivation. We find that lysosomal membrane damage elicits a robust, ubiquitin-dependent assembly of p62 puncta in both neurons and astrocytes. By contrast, metabolic stress induced by nutrient deprivation stimulates the formation of p62-ubiquitin structures preferentially in neurons as compared with astrocytes. Furthermore, pharmacological reduction in ubiquitination selectively impairs p62 clustering during starvation in neurons as compared with astrocytes. These findings indicate that neurons and astrocytes differentially regulate p62-ubiquitin conjugates in a stress-dependent manner, revealing a layer of mechanistic diversity in how neural cell types engage quality control pathways.

## MATERIALS and METHODS

### Reagents

Primary antibodies for immunofluorescence include mouse anti-βIII Tubulin (R&D Systems, MAB1195), rabbit anti-AQP4 (Millipore Sigma, HPA014784), chicken anti-GFP (Aves Labs, Inc., GFP-1020), rabbit anti-p62 (Abcam, Ab56416), mouse anti-Ubiquitin (Enzo Life Sciences, ENZ-ABS840; Fig. 1-3, S3), mouse anti-Ubiquitin (Enzo Life Sciences, BML-PW8810; Fig. 3, 4, S3), and rat anti-LAMP1 (Abcam, ab25246). We used two ubiquitin antibodies because BML-PW8810 was discontinued during the course of this study, and was replaced with ENZ-ABS840. We previously validated both antibodies to specifically recognize ubiquitin [33]. Hoechst 33342 was purchased from Thermo Fisher Scientific/Molecular Probes (H3570). Secondary antibodies for immunofluorescence include goat anti-chicken Alexa Fluor 488 (Jackson ImmunoResearch Laboratories, 103-545-155), goat anti-rabbit Alexa Fluor 594 (Invitrogen, A11037), goat anti-mouse Alexa Fluor 647 (Invitrogen, A32728), and goat anti-rat Alexa Fluor 647 (Invitrogen, A21247). Small-molecules include L-Leucyl-L-Leucine methyl ester (LLOMe; Cayman Chemicals, CAYM-16008) and TAK-243 (MLN7243; SelleckChem, S8341); small molecules were dissolved in DMSO (Sigma, 472301). Earle’s balanced salt solution (EBSS) was purchased from Sigma (E3024).

**Figure 1.**
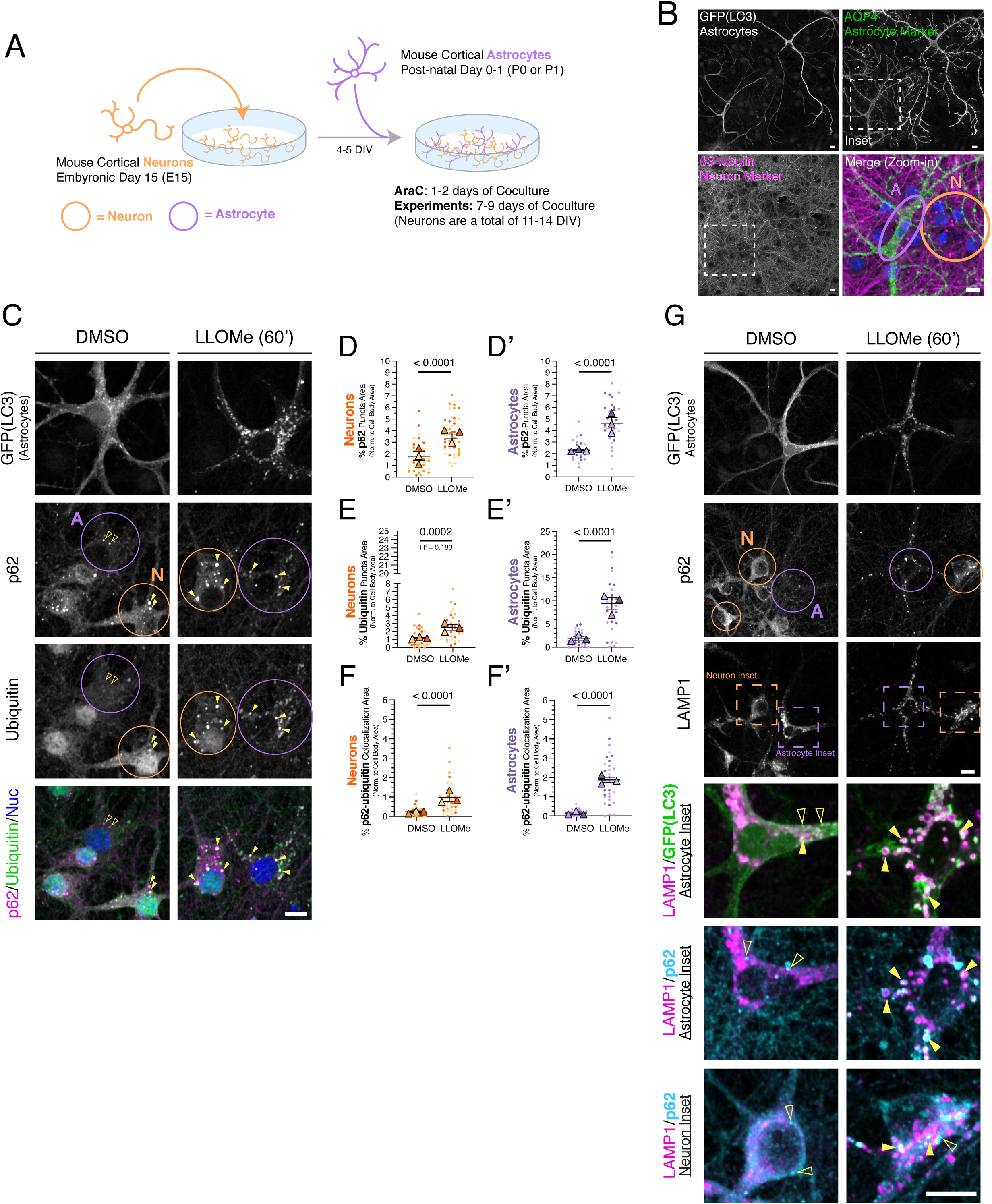
Lysosomal damage triggers the formation of p62-ubiquitin structures in both neurons and astrocytes. **(A)** Schematic of the neuron-astrocyte coculture**. (B)** Representative images of the neuron-astrocyte coculture. Coculture of GFP-LC3 transgenic astrocytes and non-transgenic neurons were fixed on DIV7 of coculture (neurons were a total of DIV11) and immunostained for GFP (LC3; labels GFP-LC3 transgenic astrocytes), AQP4 (astrocyte marker) and β3-tubulin (neuron marker); nuclei were labeled with Hoechst. Shown are maximum projections of z-stacks. Scale bars, 10 µm. **(C)** Cocultured neurons and astrocytes were treated for 1 hr with 1 mM LLOMe to induce lysosomal damage or an equivalent volume of DMSO solvent as a control. Cocultures were immunostained for GFP (LC3; labels GFP-LC3 transgenic astrocytes), p62, and ubiquitin; nuclei were labeled with Hoechst. Shown are maximum projections of z-stacks. Images for p62 and ubiquitin are grayscale-matched within a protein marker, across treatment conditions. Filled yellow arrowheads indicate colocalization between p62 puncta and ubiquitin puncta. Empty yellow arrowheads indicate p62 puncta with no ubiquitin puncta correlate. Scale bar, 10 µm. **(D-D’)** Quantification of the total area occupied by p62-positive puncta normalized to soma area for neurons **(D)** or astrocytes **(D’)**. Horizontal bars represent the means of the biological replicates ± SEM; shown are p-values from a LME model; N=37-40 neurons and N=27-33 astrocytes from 3 independent experiments; 7-8 DIV of coculture. **(E-E’)** Quantification of the total area occupied by Ub-positive puncta normalized to soma area for neurons **(E)** or astrocytes **(E’)**. Horizontal bars represent the means of the biological replicates ± SEM; shown are p-values from a LME model; N=32-33 neurons and N=27-31 astrocytes from 3 independent experiments; 7-8 DIV of coculture. **(F-F’)** Quantification of the percentage of overlapping area between p62-positive puncta and ubiquitin-positive puncta normalized to soma area for neurons **(F)** or astrocytes **(F’)**. Horizontal bars represent the means of the biological replicates ± SEM; shown are p-values from a LME model; N=31-33 neurons and N=24-31 astrocytes from 3 independent experiments; 7-8 DIV of coculture. **(G)** Maximum projections of cocultured neurons and astrocytes immunostained for p62, LAMP1, and GFP (LC3; labels GFP-LC3 transgenic astrocytes). Images for GFP(LC3), p62, and LAMP1 are grayscale-matched within a protein marker, across treatment conditions. Filled yellow arrowheads indicate puncta co-positive for LAMP1 and GFP(LC3), or LAMP1 and p62. Scale bar, 10 µm. Throughout the figure, neurons are circled in orange and astrocytes are circled in purple. For all graphs in all figures, small circles indicate the measurements from individual cells (e.g., the technical replicates) from each of the independent experiments; large triangles indicate the corresponding biological means from each of the independent experiments (e.g., the biological replicates); independent experiments are color-coded.

### Coculture of primary mouse cortical neurons and cortical astrocytes

#### Primary Cortical Neuron Culture

Transgenic mice expressing GFP-LC3 (EGFP fused to the N-terminus of rat LC3B) were obtained from the RIKEN BioResource Research Center (RBRC00806; strain B6.Cg-Tg(CAG-EGFP/LC3)53 Nmi/NmiRbrc; GFP-LC3#53) and maintained as heterozygotes. All animal protocols were approved by the Institutional Animal Care and Use Committee at the University of Pennsylvania (Protocol #805934). For procedures isolating embryos, timed-pregnant females were euthanized at E15.5 by overdose of isoflurane; euthanasia of the mother was confirmed by cervical dislocation and euthanasia of the embryos was confirmed by decapitation. Cerebral cortices were dissected from brains of GFP-LC3 transgenic mouse embryos (or non-transgenic littermates) of either sex at embryonic day 15.5. GFP-LC3 transgenic brain tissue was distinguished from non-transgenic brain tissue at the start of the dissection based on the presence or absence of GFP fluorescence, respectively. Tissue was then digested and cortical neurons were isolated following our published protocols [32, 35]. Neurons were plated at 3 million cells per 10-cm dish containing eight 25-mm acid-washed glass coverslips, or 1.15 million cells per 6-cm dish containing twelve 12-mm coverslips or five 18-mm coverslips (plating density of ∼49,000 cells/cm^2^), coated with 0.1 mg/mL poly-L-lysine (Sigma, P2636). Neurons were plated in attachment media (MEM [11095-072], 10% (v/v) heat-inactivated horse serum [Gibco, 16050-122], 1 mM sodium pyruvate [Gibco, 11306-070], 33 mM glucose [Sigma,G8769], and 37.5 mM NaCl) and incubated at 37°C in a 5% CO_2_ incubator for 1-4 hours. In Figure 4, GFP-LC3 transgenic neurons were diluted 1:10 with non-transgenic neurons and plated onto 10 cm-dishes containing 8 x 25 mm coverslips at the density described above. Neurons were then transferred into either 12-well dishes (for 12-mm coverslips) or 6-well dishes (for 18-mm or 25-mm coverslips) and maintained for 4-5 DIV in neuron maintenance media (Neurobasal medium [Gibco, 21103-049] supplemented with 2% B-27 [Gibco,17504–044], 37.5 mM NaCl, 33 mM glucose [Sigma,G8769], 2 mM glutaMAX, and 100 U/ml penicillin and 100 μg/ml streptomycin) at 37°C in a 5% CO_2_ incubator before the addition of astrocytes (discussed below). In some experiments in Figure 3 and S3, neurons were fed on 2-3 DIV of the neuron culture, where 25% media was replaced, and 1 μM AraC (antimitotic drug; Sigma, C6645) was added to reduce glial growth.

#### Primary Cortical Astrocyte Culture

Cerebral cortices were dissected from brains of GFP-LC3 transgenic neonatal mice (or non-transgenic littermates) of either sex at post-natal day 0 or 1 (P0-P1); mouse neonates at stage P0-P1 were euthanized by decapitation. GFP-LC3 transgenic brain tissue was distinguished from non-transgenic brain tissue at the start of the dissection based on the presence or absence of GFP fluorescence, respectively. Tissue was then digested and cortical glia were isolated following our published protocols that yield glial preparations that are highly enriched for astrocytes [32, 35]. Glia were plated at 1-3 million cells per 10-cm dish and cultured in glial media (DMEM [Gibco, 11965–084] supplemented with 10% (v/v) heat inactivated fetal bovine serum [Hyclone, SH30071.03], 2 mM Glutamax [Gibco, 35050–061], 100 U/ml penicillin and 100 μg/ml streptomycin [Gibco, 15140–122]) at 37°C in a 5% CO_2_ incubator. The day after plating, and every 3-4 days following, glia were fed by fully replacing the glial media. When glia reached 70-80% confluency, astrocytes were dissociated from the 10 cm tissue culture dish using 0.05% trypsin-EDTA (Gibco, 15400-054) and plated onto neurons. Occasionally, to coordinate timing with the neuron preparation, astrocytes were passaged an additional time before plating onto neurons. To reduce microglial contamination, glial culture dishes were vigorously tapped to dislodge microglia before aspirating and discarding media when feeding or passaging cells.

#### Coculture of Primary Cortical Neurons and Astrocytes

On 4-5 DIV of neuronal culture, astrocytes were trypsinized from the 10-cm plate, as described above, and used for coculture with neurons. After dissociation from the 10-cm tissue culture dish, trypsin was inactivated using glial media. Astrocytes were then spun down at 2000 rpm for 2 minutes and resuspended in coculture media (Neurobasal supplemented with 2% B-27, 1% G-5 [Gibco, 17503-012], 0.25% GlutaMAX, 100 U/ml penicillin, and 100 µg/ml streptomycin); the G5 supplement promotes astrocyte growth and maturation. Neuronal maintenance media was removed from the wells containing neurons, and astrocytes were plated onto the neurons at a density of 5,200 or 7,000 cells/cm^2^ (depending on glial growth rate prior to glial passaging) in coculture media, totaling 50,000-67,500 astrocytes per 6 well or 20,000-25,000 glia per 12 well. The coculture was maintained at 37°C in a 5% CO_2_ incubator. After 1-2 DIV of coculture, AraC was added to the cells in 100 µl of coculture medium to reach a final concentration of 2 μM AraC to prevent cell division and promote maturation of the astrocytes. The coculture was then maintained in the incubator until experiments were performed on 7-8 DIV of coculture (11-13 DIV of neuron culture) (Fig. 1 and S1).

### Stress Treatments

#### Lysosomal damage

Cocultures were treated with 1 mM L-Leucyl-L-Leucine methyl ester (i.e., LLOMe) for 1 hr at 37°C in a 5% CO_2_ incubator. To perform the treatment, conditioned coculture media (i.e., media already present on the cells) was diluted 1:1 with fresh coculture media (pre-warmed to 37°C) supplemented with 1 mM LLOMe or an equivalent volume of DMSO as a solvent control (DMSO volume is 0.4% of total treatment volume).

#### Starvation (metabolic stress) treatment

Coculture media was replaced with either Earle’s Balanced Salt Solution (EBSS, Sigma, E3024), to starve the cells, or replaced with fresh coculture media (e.g., fed controls); both medias were pre-warmed to 37°C in a water bath prior to treatment. Treatments contained 0.1% DMSO. For short-term treatments, samples were incubated for 30 minutes at 37°C in a 5% CO_2_ incubator. For long-term treatments, samples were incubated for 4 hours at 37°C in a 5% CO_2_ incubator.

#### Inhibition of the E1 ubiquitin activating enzyme

Cocultures were pretreated for 1.5 hr with 1 µM TAK-243 (MLN7243) or an equivalent volume of DMSO as a solvent control at 37°C in a 5% CO_2_ incubator. To perform the pretreatment, conditioned coculture media was diluted 1:1 with fresh coculture media (pre-warmed at 37°C in a water bath) supplemented with 1 µM TAK-243 or an equivalent volume of DMSO as a solvent control (DMSO volume is 0.1% of total treatment volume); samples were incubated for 1.5 hr at 37°C in a 5% CO_2_ incubator. For Figure 2, after the 1.5-hour pretreatment, the pretreatment media was diluted 1:1 with fresh coculture media (pre-warmed at 37°C in a water bath) and supplemented with 1 mM LLOMe (or equivalent volume of DMSO) and TAK-243 (or equivalent volume of DMSO) to a final concentration of 1 µM TAK-243 (DMSO volume is 0.5% of total treatment volume). Samples were incubated for 1 hr at 37°C in a 5% CO_2_ incubator (Fig. 2A). For Fig. 4, after the 1.5 hr TAK-243 pretreatment, all treatment media was removed and replaced with EBSS supplemented with 1 µM TAK-243 (or DMSO as a solvent control); samples were incubated for 30 min at 37°C in a 5% CO_2_ incubator.

**Figure 2.**
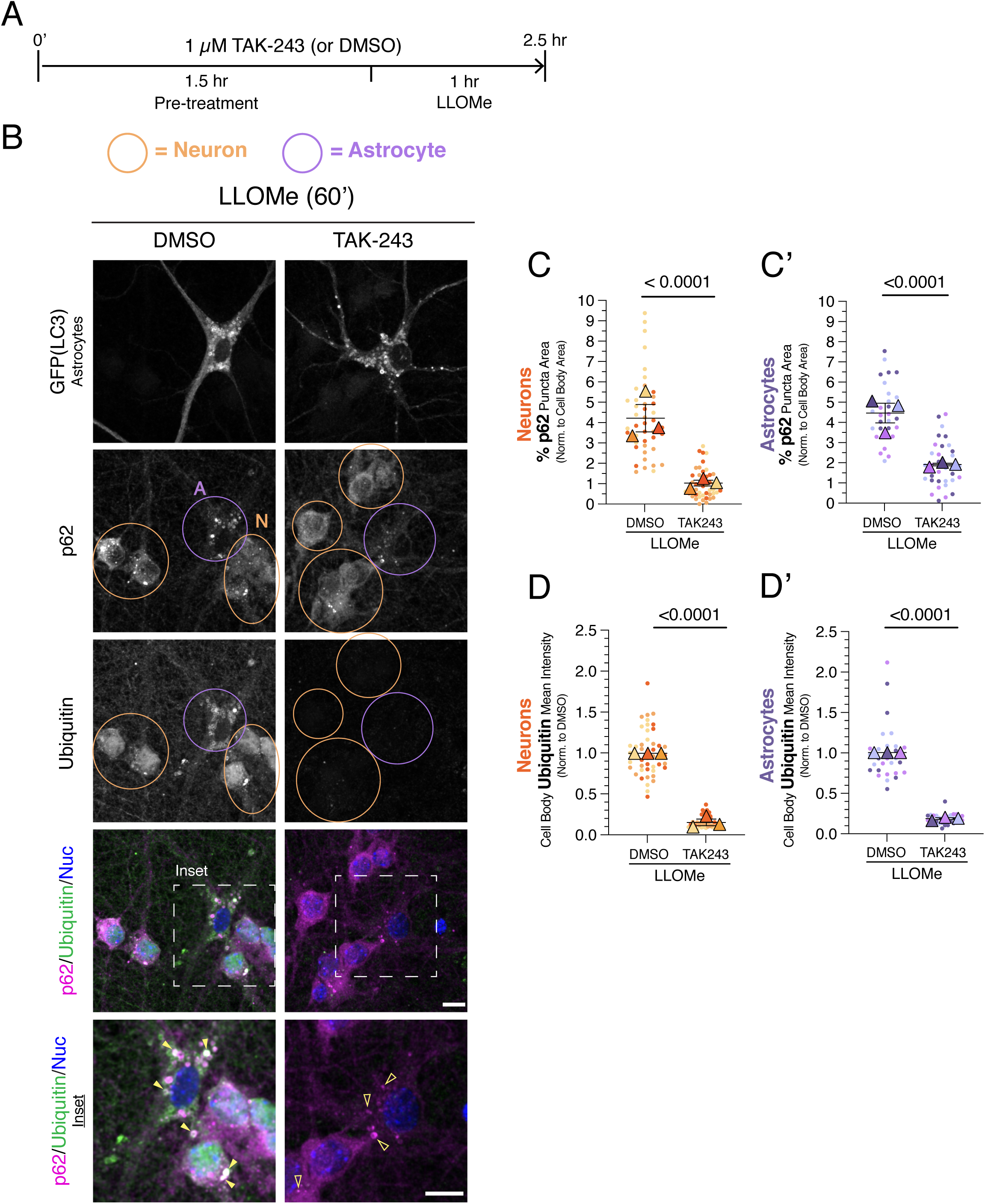
Reducing ubiquitination decreases p62-positive puncta formation during lysosomal damage in both neurons and astrocytes. **(A)** Timeline for experimental treatments. Cocultures of neurons and astrocytes were pretreated for 1.5 hr with 1 µM TAK-243 to inhibit the E1 ubiquitin-activating enzyme or an equivalent volume of DMSO solvent as a control. Cells were then incubated for 1 hour in 1 mM LLOMe to induce lysosome damage, in the continued presence of 1 µM TAK-243 or DMSO. **(B)** Immunostain analysis of co-cultured neurons and astrocytes for GFP (LC3; labels GFP-LC3 transgenic astrocytes), p62, and ubiquitin; nuclei were labeled with Hoechst. Shown are maximum projections of z-stacks. Images for p62 and ubiquitin are grayscale-matched within a protein marker, across treatment conditions. Filled yellow arrowheads indicate colocalization between p62 puncta and ubiquitin puncta. Empty yellow arrowheads indicate p62 puncta with no ubiquitin puncta correlate. Neurons are circled in orange and astrocytes are circled in purple. Scale bar, 10 µm. **(C-C’)** Corresponding quantification of total area occupied by p62-positive puncta normalized to soma area for neurons **(C)** or astrocytes **(C’)**. Horizontal bars represent the means of the biological replicates ± SEM; shown are p-values from a LME model; N=46-47 neurons and N=29-33 astrocytes from 3 independent experiments; 7-8 DIV of coculture. **(D-D’)** Quantification of the normalized mean intensity of ubiquitin signal in the cell body of neurons **(D)** and astrocytes **(D’)**. Horizontal bars represent the means of the biological replicates ± SEM; shown are p-values from a LME model; N=44-45 neurons and N=29-34 astrocytes from 3 independent experiments; 7-8 DIV of coculture.

### Immunostaining

Following treatments, cocultured cells were fixed for 10 min in 4% PFA/4% sucrose (PFA, Sigma, P6148; D-sucrose, Fisher Scientific, BP220-1) in 1X PBS (150 mM NaCl, 50 mM NaPO_4_, pH 7.4); fixative was pre-warmed to 37°C prior to cell fixation. Cells were washed three times in 1X PBS, and stored in PBS at 4°C for 1-2 days until subsequent processing. Cells were then permeabilized for 5 min in 0.1% Triton X-100 (Thermo Fisher Scientific, BP151-100) in 1X PBS, washed twice with 1X PBS, and blocked for 1 hr in 1X PBS supplemented with 5% (v/v) goat serum (Sigma, G9023) and 1% (w/v) BSA (Thermo Fisher Scientific, BP1605-100). Samples were labeled with primary antibodies diluted in blocking solution for 1 hr at room temperature. Cells were washed 3 times for 5 min each in 1X PBS, and then incubated in secondary antibody diluted in blocking solution for 1 hr at room temperature. Coculture samples were washed 3 times for 5 min each in 1X PBS, washed once in Milli-Q water, and mounted on glass microscope slides (ThermoFisher 12-544-2) in ProLong Gold (Thermo Fisher Scientific/Molecular Probes, P36930). When indicated in the figure, Hoechst (Thermo Fisher Scientific/Molecular Probes, H3570) was included at 0.5 μg/ml either during the secondary antibody incubation or in the final PBS wash before mounting coverslips. All steps of the immunostain procedure were performed at room temperature with the samples protected from ambient light.

Samples were imaged on a BioVision spinning disk confocal system consisting of a Yokagawa W1 spinning disk confocal and a Photometrics Prime 95B sCMOS camera. Images were acquired with VisiView software using either a 40X/1.4 NA (Fig. 1B) or 63X/1.4 NA (all other figures) Plan-Apochromat oil-immersion objective with either a 1X (Fig. 1-3, S2A-E, S3, S4) or 2X (Fig 4, S2F, G) optical magnifier and solid-state 405, 488, 561, and 640 nm lasers for excitation. Z-stacks were obtained that spanned the entire depth of the cell at 0.2-μm sections. Images that would be quantitatively compared to each other were obtained with the same acquisition parameters across treatment conditions and biological replicates.

**Figure 3.**
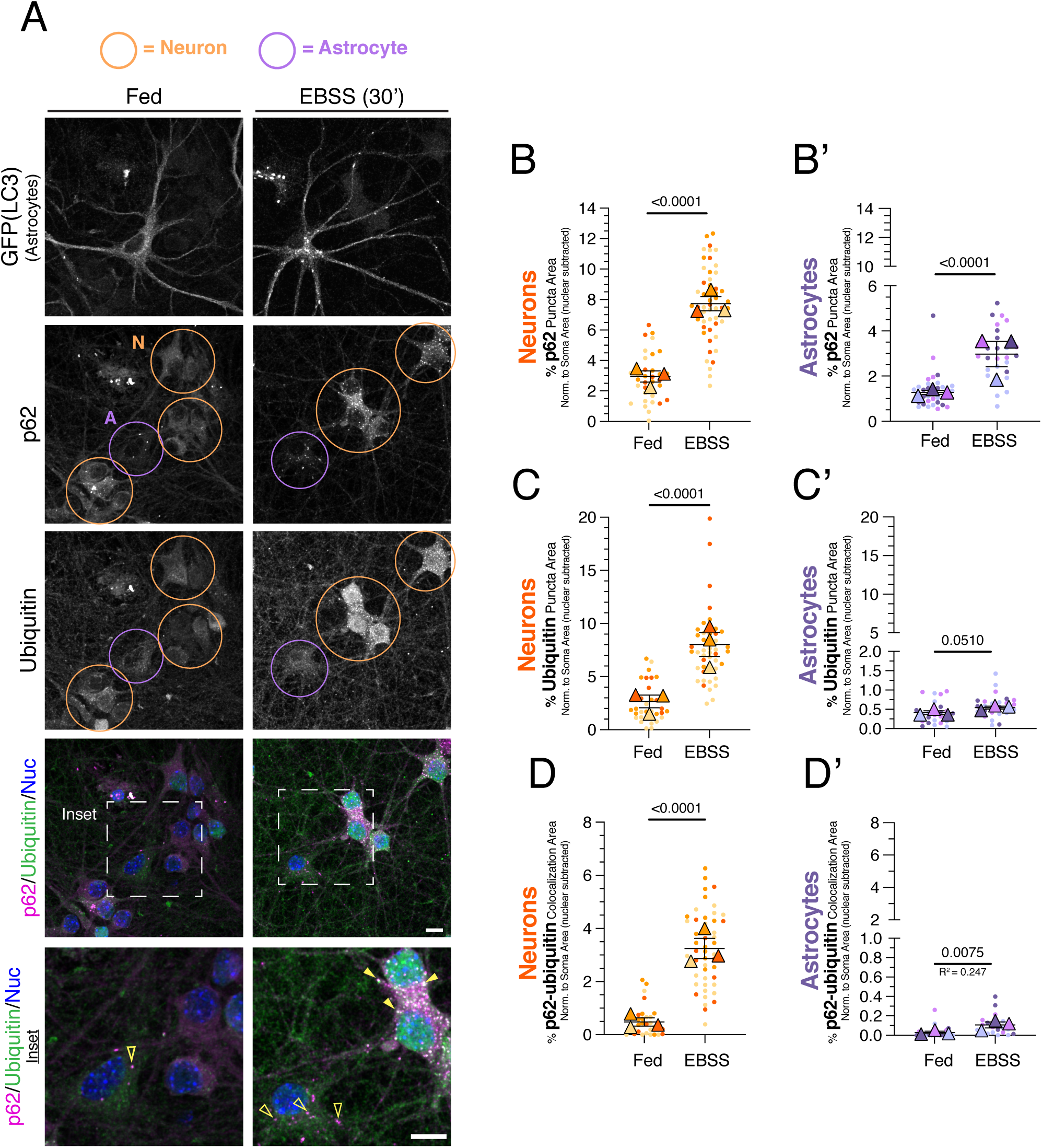
Short-term metabolic stress preferentially increases p62-Ub structures in neurons as compared to astrocytes. **(A)** Cocultures of neurons and astrocytes were treated for 30 minutes in EBSS (or regular media as a fed control) and then immunostained for GFP (LC3; labels GFP-LC3 transgenic astrocytes), p62, and ubiquitin; nuclei were labeled with Hoechst. Shown are maximum projections of z-stacks. Filled yellow arrowheads indicate colocalization between p62 puncta and ubiquitin puncta. Empty yellow arrowheads indicate p62 puncta with no ubiquitin puncta correlate. Neurons are circled in orange and astrocytes are circled in purple. Scale bar, 10 µm. **(B-B’)** Corresponding quantification of total area occupied by p62-positive puncta normalized to cytoplasmic area in the soma for neurons **(B)** or astrocytes **(B’)**. Horizontal bars represent the means of the biological replicates ± SEM; shown are p-values from a LME model; N=34-52 neurons and N=27-31 astrocytes from 3 independent experiments; 7 DIV of coculture. **(C-C’)** Corresponding quantification of area occupied by Ub-positive puncta normalized to cytoplasmic area in the soma for neurons or astrocytes. Horizontal bars represent the means of the biological replicates ± SEM; shown are p-values from a LME model; N=31-48 neurons and N=24-25 astrocytes from 3 independent experiments; 7 DIV of coculture. **(D-D’)** Quantification of the percentage of overlapping area between p62-positive puncta and ubiquitin-positive puncta normalized to cytoplasmic soma area for neurons **(D)** or astrocytes **(D’)**. Horizontal bars represent the means of the biological replicates ± SEM; shown are p-values from a LME model; N=29-48 neurons and N=22-23 astrocytes from 3 independent experiments; 7 DIV of coculture.

**Figure 4.**
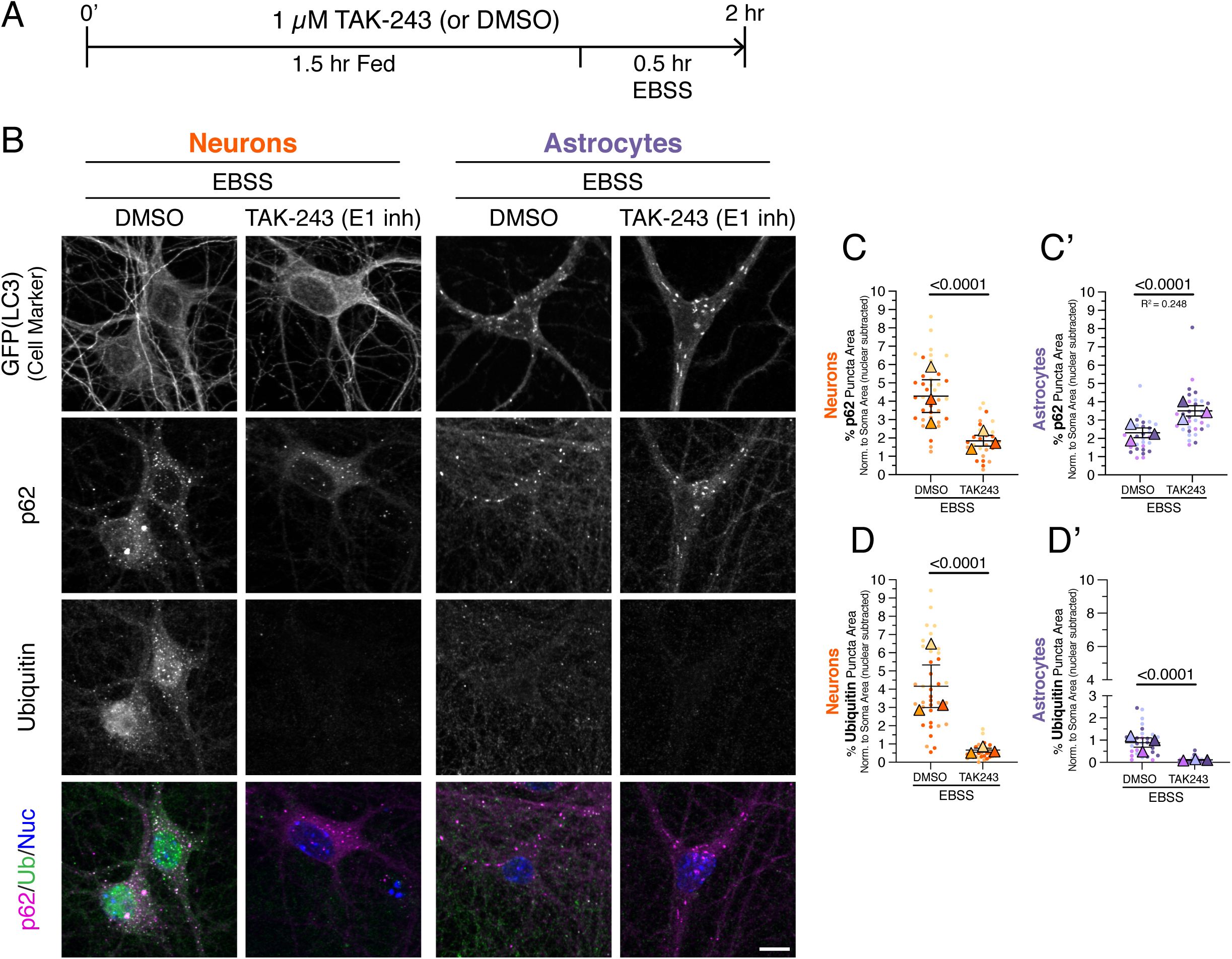
Reducing ubiquitination decreases p62 puncta formation during metabolic stress in neurons but not in astrocytes. **(A)** Timeline for experimental treatments. Cocultures of neurons and astrocytes were pretreated for 1.5 hr with 1 µM TAK-243 to inhibit the E1 ubiquitin activating enzyme or an equivalent volume of DMSO solvent as a control. Cells were then incubated for 30 minutes in EBSS to induce nutrient deprivation, in the presence of 1 µM TAK-243 or DMSO. **(B)** Immunostain analysis of cocultured neurons and astrocytes for GFP(LC3), p62, and ubiquitin; shown are maximum projections of z-stacks. To identify neurons, GFP-LC3 transgenic neurons were cocultured with non-transgenic astrocytes. Since lipidated GFP-LC3 is known to not preserve well during fixation specifically in neurons, the GFP-LC3 transgene serves only as a cell-type-specific marker to identify neurons. To identify astrocytes, GFP-LC3 transgenic astrocytes were cocultured with non-transgenic neurons. Nuclei were labeled with Hoechst. Scale bar, 10 µm. **(C-C’)** Quantitation of total area occupied by p62-positive puncta normalized to cytoplasmic soma area for neurons **(C)** or astrocytes **(C’)**. Horizontal bars represent the means of the biological replicates ± SEM; shown are p-values from a LME model; N=30-40 neurons and N=32-34 astrocytes from 3 independent experiments; 7-8 DIV of coculture. **(D-D’)** Quantitation of total area occupied by Ub-positive puncta normalized to cytoplasmic soma area for neurons **(D)** or astrocytes **(D’)**. Horizontal bars represent the means of the biological replicates ± SEM; shown are p-values from a LME model; N=30-38 neurons and N=32-34 astrocytes from 3 independent experiments; 7-8 DIV of coculture.

### Image Analysis

#### Area occupied by GFP(LC3), p62, and ubiquitin positive puncta

We first generated maximum projections of Z-stacks in FIJI. Astrocytes were selected for analysis based on morphological complexity, containing at least 3 well-defined processes that developed secondary and tertiary branches. Neurons were selected for analysis based on the following criteria: neurons must exhibit a well-defined axon and at least 3 well-defined and branching dendrites, one of which being a primary dendrite. For Fig. 4 and S4, astrocyte and neuronal somas were outlined using the GFP(LC3) channel and measured for total cross-sectional area; the ROIs outlining the somas were saved for subsequent use. For all other figures, neurons were selected and outlined in the p62 channel, using the cytosolic levels of p62 as a space fill, and astrocytes were selected and outlined using the GFP-LC3 channel. ROIs outlining the somas were saved for subsequent use. We then removed signal outside the ROI using the FIJI function “clear outside” and the individual cell images were then loaded into Ilastik to identify and segment puncta of interest. To reduce bias in selecting cells for analysis, images were blinded to treatment identity during cell selection using the “Filename Randomizer” plugin (credit: Tiago Ferreira) for the following figures: Fig. 3, S2B-E, and S3. Ilastik was used to identify and segment GFP(LC3)-positive puncta (as measured using the GFP antibody), p62-positive puncta, and ubiquitin-positive puncta. To account for cell-type-specific differences in signal intensity, we trained separate Ilastik programs for each marker across treatment paradigms for each cell type. The only exception is that we used the same Ilastik program for quantifying ubiquitin puncta in astrocytes for both the 30 minutes and 4 hours EBSS treatment paradigms (as in Fig. 3C’ and S3C’). To enable direct comparisons, the same Ilastik program was always used for all replicates of a given experiment. Ilastik segmentations were then imported into FIJI to generate a mask of the segmentation. The total area occupied by segmented puncta was then measured using the “analyze particles” function in FIJI. Total area occupied by LC3, p62, and ubiquitin-positive puncta per cell was normalized to the corresponding cell body area (or cytoplasmic area, as described below) and plotted as a percentage of total cell body area. To calculate the average size of puncta (e.g., Fig. S4), the total puncta area was divided by the number of puncta per cell; values obtained using the “analyze particles” function in FIJI.

#### Nuclear subtraction

In the starvation treatments, we observed that ubiquitin puncta increased in the nucleus of neurons. This observation may be related to known functions for ubiquitin in regulating proteasomal activity and histone modification to regulate DNA accessibility in certain paradigms of stress [36–39]. Since we were primarily interested in quantifying the population of ubiquitin in the cytoplasm, we removed the nuclear area from the cell body area (Fig. 3, 4, S2B-S2G, S3, S4). To do this, we first trained an Ilastik program to recognize the nucleus based on the Hoechst stain for each outlined cell. We then created a mask of the nuclear signal in FIJI and inverted the mask to create a mask of the cytoplasm. Using the “AND” function in the image calculator tool in FIJI, we counted pixels that were present in both the cytoplasm mask and the mask of the total puncta for each protein of interest. The area of cytoplasmic puncta for each protein was then normalized to the cytoplasmic area for each individual cell and expressed as a percentage.

#### Co-localization area

To measure colocalization between p62 and GFP(LC3) or ubiquitin puncta, we determined the overlapping area between the Ilastik segmentations for p62, GFP(LC3), and ubiquitin puncta that were generated as described above. Using FIJI, we then measured the overlapping area between indicated markers using the “AND” function in the image calculator tool for each individual cell. For starvation datasets, we used the nuclear-subtracted segmentations for colocalization analysis. Total overlapping area was then normalized to the corresponding soma area (or cytoplasmic area for nuclear-subtracted analysis) and expressed as a percentage.

#### Soma Ubiquitin analysis

For Fig. 2D-D’, we used the ROIs of the soma outlines generated above to measure the ubiquitin signal in the soma. The ROI outlining the soma was overlaid onto the maximum projection of the ubiquitin channel for each individual cell, and we measured the mean gray value of the ubiquitin signal using FIJI. For each condition and biological replicate, we also measured the mean gray value of a circular ROI that was drawn in an area with no cellular processes or meshwork to represent the background signal. This mean gray value for background was then subtracted from the corresponding ubiquitin mean gray value for each individual cell to create a background-corrected mean gray value for ubiquitin (hereafter referred to as “mean gray value”). We then normalized the mean gray values for each individual cell in both the control and treatment groups to the corresponding DMSO control average for each independent experiment and each cell type. Normalized mean gray values for each soma (e.g., technical replicates) were averaged to generate a mean for each independent experiment (e.g. biological replicates) and plotted in PRISM.

### Figure and Manuscript Preparation

For presentation of images, maximum and minimum gray values were adjusted linearly in FIJI. Images that are grayscale matched are denoted in the relevant figure legend. Superplots of graphs were assembled in Graphpad Prism according to published techniques [40]. In all graphs in all figures, small circles indicate the measurements from individual cells (e.g., the technical replicates) from each of the independent experiments; large triangles indicate the corresponding biological means from each of the independent experiments (e.g., the biological replicates); bars are the means of the biological means ± SEM; independent experiments are color-coded. Figures were assembled using Adobe Illustrator. The microscope image in Figure S1 was generated using the “Generate Vector” AI tool in Illustrator. Perplexity and ChatGPT AI tools were used to make some sentences and paragraphs more concise to improve clarity and readability. After use of these tools, the authors carefully reviewed and edited the content as needed and take full responsibility for the content of the publication.

### Statistical analyses

All image measurements were obtained from the raw data. For all datasets, we used RStudio (Version 2025.5.0.496) to run a linear mixed effects model (LME, R package “lme4”). Using a LME model enables us to account for variation in the data that is unrelated to the treatment condition (modeled as random effects), a feature absent in standard ANOVA models. A LME also recognizes that cells originating from the same treatment group (e.g., cells on the same coverslip) are not independent, reducing the risk of inflating statistical power. We assigned treatment condition (e.g. LLOMe or TAK-243) as the fixed effect, and the biological replicate (e.g. cells from each independent preparation from a different mouse) as the random effect. To confirm that the model fit was appropriate, we assessed the homoscedasticity of the model (i.e., whether the variance of the data is equal across the entire range of the fixed effect) for each dataset analyzed. In datasets where the model was not homoscedastic, we utilized the “VarIdent” function (R package “nlme”) to standardize the variance between different treatment conditions and improve the LME model fit. We used the “VarIdent” function for the following datasets: Figures 1E’, 3B’, 3D, 3D’, 4D, S2C-S2E, S3B, S3B’, S3D, S4B. All other datasets used the unadjusted LME model. To confirm that the results of the LME model did not differ between the two LME packages used (lme4 versus nlme), we ran several LME models (without any corrections) using the same data in both packages and found the p-values to be nearly identical. We used marginal R^2^ values to assess how much of the variance was explained by the model. R^2^ values of 0.1-0.25 are considered modest effects, whereas R^2^ values less than 0.1 are considered weak effects. Unless noted directly in the figure, R^2^ values were above a cutoff of 0.25, indicating that the treatment variable (e.g., LLOMe) accounts for a substantial portion of the variance. When an R^2^ value was below 0.25 but statistical significance was achieved, this combination indicates that the treatment variable has a modest effect, but explains only a limited amount of the outcome variance and is noted in the relevant graph. When an R^2^ value was below 0.1 but statistical significance was achieved, this combination indicates that the treatment variable has a weak effect, but explains only a very limited amount of the outcome variance and is noted in the relevant graph. All quantifications were performed with 3 independent experiments, and comparisons were only made within cell type, not between cell type. P-values are denoted directly within each graph.

## RESULTS

### Neurons and astrocytes exhibit similar p62-ubiquitin responses to lysosomal damage

To investigate the regulation of p62-ubiquitin conjugates in neurons and astrocytes in models of cellular stress, we cocultured primary mouse cortical neurons with primary mouse cortical astrocytes. Neurons (∼96% pure based on our prior analysis with a neuron-specific marker [32]) are cultured for 4-5 days *in vitro* (DIV) to establish a dense meshwork of neurites before addition of astrocytes (Fig. 1A, S1). Astrocytes were then plated on top of the neuron meshwork and cocultured for 7-8 DIV, resulting in a total neuronal age of 11-13 DIV; Fig 1A, S1). This order of addition is critical as direct contact and secreted factors from developed neurons promote astrocyte maturation [41, 42]. In this context, astrocytes develop arborized processes reminiscent of morphologies observed *in vivo* (Fig. 1B). Immunostaining for aquaporin 4 (AQP4), an astrocyte-enriched water channel, and β3-tubulin, a marker of neuronal microtubules, reveals the branched morphologies of astrocytes in coculture with a dense neuronal network (Fig. 1B).

We also made use of a transgenic mouse expressing GFP-LC3 [43] to distinguish cell-types (Fig. S1). In its cytosolic form, LC3 can delineate cellular morphologies and boundaries and, when lipidated, LC3 labels autophagosome membranes. This tool enables visualization of morphological compartments, quantification of autophagy levels, and validation of stress-inducing treatments. To distinguish astrocytes from neurons, astrocytes from GFP-LC3 transgenic mice were cocultured with non-transgenic neurons (Fig. 1B, S1C). Based on our previously published purity analysis [32], all non-transgenic cells with neuronal morphology were classified as neurons (see Methods). Together, this neuron-astrocyte coculture provides a method to examine cell-type-specific regulation of p62-ubiquitin conjugates in response to defined stress paradigms.

We first examined cell-type-specific responses to lysosomal membrane damage elicited by L-leucyl-L-leucine methyl ester (LLOMe), a lysosomotropic agent that is processed by cathepsin C to generate membranolytic polymers that permeabilize lysosomal membranes [44]. Sustained exposure to LLOMe promotes the autophagic removal of damaged lysosomes, a process known as lysophagy, which involves the recruitment of selective autophagy receptors, including p62, to ubiquitinated substrates on the lysosome [3, 26, 45, 46]. This pathway has been extensively characterized in immortalized cell lines, but little is known about how neurons and astrocytes respond to lysosomal injury, despite pathophysiological relevance for lysosomal quality control in neurodegenerative disease.

To induce lysosomal membrane damage, cocultures were treated with 1 mM LLOMe for 1 hr; astrocytes were identified by expression of GFP-LC3. The effectiveness of the treatment was validated by quantifying GFP-LC3-positive puncta in astrocytes. During early stages of lysosomal damage, LC3, an Atg8 family protein, is directly conjugated to lysosomal membranes in a process termed CASM (conjugation of ATG8 to single membranes), which may support lysosomal repair [47, 48]. More prolonged damage leads to LC3 recruitment during lysophagy [49]. Thus, the appearance of LC3 on lysosomes provides a readout for effective induction of lysosomal injury. To quantify GFP-LC3 puncta, we used Ilastik, a machine-learning platform trained to segment punctate structures from diffuse cytosolic signal. LLOMe treatment caused a marked increase in GFP-LC3-positive puncta and ring-like structures in astrocytes relative to the solvent control (Fig. 1C, S2A). These results confirm that LLOMe efficiently induces lysosomal damage and triggers the formation of LC3-positive structures, consistent with activation of CASM, lysophagy, or both.

Next, we immunostained neuron-astrocyte cocultures for p62 and ubiquitin. We used two ubiquitin antibodies that detect both mono- and polyubiquitinated chains, including K29-, K48-, and K63-linked species. As validated in our previous work, these antibodies recognize p62-positive fibrils and aggresomes that form in astrocytes under conditions of proteasomal inhibition [33]. In DMSO-treated controls, occasional p62-positive puncta were detected in both neurons and astrocytes (Fig. 1C), consistent with basal homeostatic functions for p62. In neurons, ubiquitin localized primarily to the nucleus with variable levels of cytoplasmic puncta (Fig. 1C). Nuclear localization is consistent with the established role of ubiquitin in chromatin remodeling [50]. In astrocytes, ubiquitin levels were lower overall and predominantly cytoplasmic (Fig. 1C).

LLOMe treatment significantly increased the formation of p62- and ubiquitin-positive puncta in the cytoplasm of both neurons and astrocytes relative to solvent controls (Fig. 1C-E’). However, we note that the increase in ubiquitin-positive puncta was more prominent in astrocytes than neurons (Fig. 1E-E’). Since the effects of LLOMe were most prominent in the somas of neurons and astrocytes, where degradative lysosomes are known to concentrate [51–54], we focused our analysis in this region in each cell type (Fig. 1C). LLOMe exposure also increased colocalization between p62 and ubiquitin-positive puncta in both neurons and astrocytes, (Fig. 1C, F-F’). A population of p62 and ubiquitin assembled into ring-like structures, which appeared more abundant in astrocytes (Fig. 1C). These structures resemble those previously reported to form during p62 engagement with ubiquitinated substrates on damaged lysosomes [3, 25, 26]. Consistent with this interpretation, p62 localization partially overlapped with LAMP1, a marker of late endosomes and lysosomes, in both neurons and astrocytes (Fig. 1G). LAMP1 also exhibited partial overlap with GFP-LC3 (Fig. 1G), further supporting recruitment of p62 to damaged lysosomes. Together, these data demonstrate that lysosomal membrane damage induces recruitment of p62 and ubiquitin to lysosomes in both neurons and astrocytes, with both cell types exhibiting similar responses.

### p62 recruitment to damaged lysosomes depends on ubiquitin in both neurons and astrocytes

Previous studies established that ubiquitination of lysosomal proteins is a key step in recruiting p62 to damaged lysosomes, work performed mostly in non-neural cell types [3, 25, 26]. To define the role of ubiquitin in p62 recruitment during lysosomal damage in neurons and astrocytes, we pharmacologically reduced ubiquitination using the E1 enzyme inhibitor TAK-243. Substrate ubiquitination proceeds through an enzymatic cascade involving E1 ubiquitin-activating enzymes, E2 ubiquitin-conjugating enzymes, and E3 ubiquitin ligases, which attach ubiquitin to lysine residues on target proteins. TAK-243 binds to the ATPase pocket of the E1 enzyme, thereby preventing transfer of ubiquitin to E2 enzymes and blocking downstream substrate ubiquitination [55].

To ensure that substrates were not sequestered within organelles (e.g., autophagosomes or lysosomes) and thus shielded from the TAK-243 treatment, cocultures were pretreated with TAK-243 prior to induction of lysosomal damage. Neuron-astrocyte cocultures were treated for 1.5 hours with 1 µM TAK-243 or an equivalent volume of DMSO as solvent control (Fig. 2A). Lysosomal damage was then induced by adding 1 mM LLOMe for 1 hour in the continued presence of 1 µM TAK-243 or DMSO, followed by immunostaining for GFP-LC3, p62, and ubiquitin (Fig. 2A).

In DMSO-pretreated controls, LLOMe triggered the formation of p62- and ubiquitin-positive puncta and ring-like structures in both neurons and astrocytes (Fig. 2B), consistent with the results shown in Fig. 1. Ubiquitin-positive rings co-labeled with p62 were particularly prominent in astrocytes (Fig. 2B), matching structures previously linked to lysosomal damage. Pretreatment with TAK-243 significantly reduced ubiquitin-positive structures induced by LLOMe in both neurons and astrocytes (Fig. 2B, D, D’), confirming effective inhibition of ubiquitination and validating antibody specificity. TAK-243 pretreatment also decreased the formation of p62-positive puncta in both cell types (Fig. 2B-C’), indicating that ubiquitin likely contributes to the formation of p62 structures upon lysosomal damage in neurons and astrocytes.

Given that LC3 is a ubiquitin-like protein, we verified that TAK-243 treatment did not impair LC3 lipidation. Indeed, LLOMe effectively induced the formation of LC3-positive structures in the presence of TAK-243, indicating that LC3 conjugation was not impeded by TAK-243 (Fig. 2B). Together, these results demonstrate that neurons and astrocytes similarly depend on ubiquitin for recruitment of p62 to damaged lysosomes. This finding demonstrates that mechanisms previously defined mostly in immortalized cell lines are conserved in neurons and astrocytes. Furthermore, these data validate our experimental tools for probing ubiquitin-dependent regulation of p62 under other stress paradigms.

### Neurons and astrocytes distinctly regulate p62-ubiquitin conjugates during metabolic stress

We next investigated whether neurons and astrocytes distinctly regulate p62-ubiquitin conjugates in response to metabolic stress. Thus, we examined the regulation of these conjugates under nutrient deprivation. During nutrient deprivation, autophagy is activated to degrade cellular components to supply amino acids as biosynthetic fuel for maintaining essential functions [32, 56, 57]. Nutrient deprivation can also increase p62-ubiquitin conjugates in HEK293 cells [24], but whether this result extends to neurons and astrocytes is unknown. Cocultures were starved in Earle’s Balanced Salt Solution (EBSS), a nutrient deprivation model lacking amino acids with reduced glucose compared to standard coculture medium (5.6 mM D-glucose versus 25 mM in standard coculture media). Cocultures were exposed to EBSS for 30 minutes to model short-term metabolic stress. We previously established that 30 minutes of EBSS treatment is sufficient to starve monocultured neurons and astrocytes, as indicated by reduced mTOR signaling [32].

After starvation, cocultures were immunostained for p62, ubiquitin, and GFP-LC3; astrocytes were identified by GFP-LC3 expression. We noted that starvation caused the formation of ubiquitin-positive puncta within the nucleus in neurons. To ensure that our quantification only captured ubiquitin structures linked to quality control in the cytoplasm, we excluded the area of the nucleus and quantified only cytoplasmic puncta within the soma. Thirty minutes of EBSS treatment significantly increased p62-positive puncta in both neurons and astrocytes relative to fed controls (Fig. 3A-B’). This clustering of p62 is consistent with engagement of cargoes that might be routed for autophagic degradation in response to metabolic stress. Strikingly, however, EBSS treatment increased ubiquitin-positive puncta more strongly in neurons than in astrocytes (Fig. 3A, C-C’). Furthermore, the degree of colocalization between p62 and ubiquitin puncta also increased more prominently in neurons (Fig. 3A, D-D’). This attenuated ubiquitin response in astrocytes contrasted with the lysosomal damage paradigm, where both neurons and astrocytes exhibited a robust increase in p62 and ubiquitin colocalization (Fig. 1, 2).

Since astrocytes exhibited an attenuated ubiquitin response to EBSS treatment, we wanted to confirm that astrocytes were effectively starved. Thus, we quantified GFP-LC3-positive puncta and overlap between LC3 and p62. Thirty minutes of EBSS treatment significantly increased both measurements (Fig. 3A, S2B-S2C). These findings confirm that astrocytes experienced nutrient deprivation and activated autophagy accordingly.

Given that short-term starvation produced distinct responses in neurons and astrocytes, we next examined whether prolonged nutrient deprivation would elicit similar differences in p62-ubiquitin regulation across these cell types. Thus, we starved cocultures in EBSS for 4 hours to model long-term metabolic stress. Consistent with short-term nutrient depletion, 4 hours of EBSS increased p62-positive puncta in both neurons and astrocytes (Fig. S3A-B’). However, ubiquitin-positive puncta again accumulated more prominently in neurons (Fig. S3A, C-C’), and colocalization between p62 and ubiquitin puncta remained greater in neurons than in astrocytes (Fig. S3A, D-D’). Extended starvation also increased GFP-LC3 puncta and LC3-p62 overlap in astrocytes (Fig. S2D-S2E), confirming activation of autophagy. Together, these findings indicate that both short- and long-term nutrient deprivation preferentially induce formation of p62-ubiquitin conjugates in neurons relative to astrocytes. Thus, neurons and astrocytes can diverge in how they deploy p62 and ubiquitin in a manner that depends on the specific stress.

### p62 clustering during metabolic stress depends more strongly on ubiquitin in neurons than in astrocytes

To define the role of ubiquitin in regulating p62 during metabolic stress, we reduced ubiquitination using TAK-243, as described in Fig. 2. Cocultures were pretreated for 1.5 hours with 1 µM TAK-243 or with DMSO as a solvent control, then subjected to EBSS starvation for 30 minutes in the continued presence of the inhibitor or control (Fig. 4A). To distinguish cell types, neurons or astrocytes derived from GFP-LC3 transgenic mice were cocultured with their non-transgenic counterpart (Fig. 4B, S1C). Consistent with earlier results, EBSS treatment in the DMSO pretreatment group increased p62 puncta in both neurons and astrocytes but enriched ubiquitin puncta preferentially in neurons (Fig. 4B).

Pretreatment with TAK-243 effectively reduced ubiquitin-positive structures in both neurons and astrocytes (Fig. 4B, D-D’). Interestingly, inhibition of ubiquitination selectively decreased p62 puncta density in neurons but not in astrocytes (Fig. 4B, C-C’). In astrocytes, TAK-243 treatment instead increased the total area occupied by p62 puncta (Fig. 4B, C’). This increase in p62 area was not due to a change in the number of p62 puncta (Fig. S4A), but might be due to an increase in the mean size of p62 puncta per cell (Fig. S4B). The percentage of p62 puncta colocalized with GFP-LC3 was unaltered with TAK-243 in astrocytes (Fig. S2G), indicating continued routing of p62 to autophagosomes. Moreover, TAK-243 did not reduce GFP-LC3 puncta in astrocytes during starvation (Fig. S2F), indicating that autophagosome formation itself was not impaired by reduced ubiquitination.

Together, these results demonstrate that neurons depend more strongly on ubiquitin to cluster p62 during metabolic stress than astrocytes. Hence, we reveal that distinct mechanisms underlie how different neural cell types use ubiquitin to regulate p62. Collectively, our work establishes a foundation for examining how differential regulation of p62-ubiquitin conjugates in neurons and astrocytes may influence selective vulnerability and proteostasis failure in neurodegenerative disease.

## DISCUSSION

Here, we examined how neurons and astrocytes differentially regulate p62-ubiquitin conjugates in response to two distinct forms of cellular stress: lysosomal membrane damage and metabolic stress induced by nutrient deprivation. To define these cell-type-specific responses, we used a neuron-astrocyte coculture system that reproduces key morphological, proteomic, and functional features of neuron-astrocyte interactions observed *in vivo*. Lysosomal membrane damage triggered the formation of punctate and ring-like structures positive for p62 and ubiquitin in both neurons and astrocytes (Fig. 1). The formation of these p62-positive structures was dependent on ubiquitin in both cell types (Fig. 2). Thus, neurons and astrocytes respond similarly to lysosomal membrane injury by engaging p62-ubiquitin conjugates, consistent with mechanisms described in immortalized cell lines where p62 functions in lysophagy.

In contrast, metabolic stress induced by amino acid deprivation and reduced glucose elicited divergent responses between neurons and astrocytes. Both short-term (30 minutes) and long-term starvation (4 hours) increased p62-ubiquitin structures more prominently in neurons than in astrocytes (Fig. 3, S3). Neurons also showed a greater reliance on ubiquitin to form p62 puncta during metabolic stress (Fig. 4). Even when ubiquitination was reduced by E1 inhibition, astrocytes efficiently delivered p62 to autophagosomes (Fig. S2G), and autophagosome formation itself was not impaired (Fig. S2F). Collectively, these results demonstrate that neurons and astrocytes differentially regulate p62 and ubiquitin in a stress-specific manner, highlighting context- and cell-type-specific control of proteostasis pathways.

A potential model emerging from these findings is that neurons and astrocytes may target distinct cargoes for degradation under metabolic stress. The strong dependence on ubiquitin in neurons suggests that they may be more selective in the cargoes routed for degradation during nutrient deprivation. In contrast, the relatively muted ubiquitin response in starved astrocytes implies a greater reliance on bulk autophagy, which can non-selectively engulf cytoplasmic material. These observations raise the possibility that neurons may engage more selective quality control mechanisms, whereas astrocytes may favor generalized clearance pathways to preserve metabolic stability.

Why would neurons and astrocytes exhibit different proteostatic responses to metabolic stress? These cell type differences may be due, in part, to their distinct yet interconnected metabolic programs. Astrocytes are highly glycolytic, whereas neurons depend primarily on oxidative phosphorylation for ATP production [58–62]. Metabolic coupling between neurons and astrocytes allows synaptic activity to stimulate glycolysis in astrocytes, leading to lactate production and export to neurons, where it serves as an energy substrate and supports long-term memory formation [58, 62–65]. Astrocytes also release ATP, which modulates neuronal activity [66–69], and lysosomes enriched in ATP have been proposed as a source of extracellular ATP released through lysosomal exocytosis [70, 71]. Extracellular ATP can be metabolized to adenosine which activates neuronal adenosine receptors to modulate neuronal activity [66–69]. In total, these different metabolic roles may require astrocytes to deploy p62-dependent homeostatic mechanisms differently than neurons. In this way, astrocytes may also use autophagy to mobilize resources to support metabolic and synaptic demands of neurons.

This cooperative metabolic relationship extends to lipid and redox homeostasis, which may also impose cell-type-selective mechanisms of quality control that implicate p62. During periods of high synaptic activity, toxic peroxidated fatty acids generated in neurons are transferred to astrocytes, stored in lipid droplets, and subsequently catabolized [72]. This process triggers an astrocytic transcriptional response that mitigates oxidative stress [72]. Additionally, synaptic activity or oxidative stress can activate the transcription factor NRF2 in astrocytes, which drives expression of antioxidant genes [73, 74]. This NRF2 response exerts neuroprotective effects, potentially through transfer of antioxidants that alleviates oxidative stress in neurons [74–78].

Intriguingly, beyond roles in selective autophagy, p62 also promotes antioxidant signaling. p62 activates NRF2 by binding to KEAP1 [15, 79], an adaptor of the Cullin-3-type ubiquitin ligase complex that targets NRF2 for proteasomal degradation [80–82]. By sequestering KEAP1, p62 disrupts the KEAP1-NRF2 interaction, allowing NRF2 to translocate to the nucleus and induce transcription of antioxidant genes [15, 79]. Oxidative stress promotes phosphorylation of p62 within the KEAP1-interacting region, which enhances affinity for KEAP1 [17]. Notably, p62 is also a transcriptional target of NRF2, forming a positive feedback loop that amplifies the antioxidant response [83]. Various forms of nutrient deprivation, including amino acid, serum, or glucose starvation, are known to elevate reactive oxygen species (ROS) and increase oxidative stress [84–86]. Moreover, a recent study demonstrates that nutrient deprivation increases p62 phosphorylation and activates NRF2-dependent antioxidant signaling [87]. Thus, metabolic stress may engage p62 as an autophagy receptor and as a modulator of redox homeostasis. Since NRF2 signaling has been reported to be enriched in astrocytes [73, 74], the p62 structures that we observed in astrocytes under metabolic stress may participate in the KEAP1-NRF2 pathway to promote antioxidant defense. In this model, metabolic stress could elicit distinct functional outcomes for p62 in neurons and astrocytes, favoring selective autophagy in neurons and antioxidant signaling in astrocytes. Furthermore, ALS-FTLD-linked mutations in p62 disrupt both selective autophagy and NRF2-mediated antioxidant pathways [88]. Such disease-associated variants may therefore impair these cell-type-specific functions of p62, creating distinct neuronal and astrocytic vulnerabilities that collectively drive neurodegenerative progression. Together, these observations highlight how stress-dependent regulation of p62 may influence selective vulnerability in ALS and related disorders. Future studies will be needed to test this model and define how p62 coordinates divergent stress responses across cell types.

Another possible mechanism for the cell-type-specific divergence in p62 regulation is that p62 structures formed in astrocytes during metabolic stress may correspond to stress granules. Stress granules assemble in response to diverse stress conditions, including nutrient deprivation, particularly under reduced or depleted glucose levels [89–93]. Several studies have shown that p62 associates with stress granules formed during arsenite-induced oxidative stress and promotes their clearance through autophagy [11, 12]. Notably, the association of p62 with stress granules occurs through multiple mechanisms, including those that may be distinct from ubiquitination. Chitiprolu et al. reported that ubiquitin is not enriched in stress granules [11], and proposed that p62 is recruited through interactions with arginine-dimethylated proteins, such as FUS, which are abundant in these structures [11]. In addition, Jeon et al. found that p62-mediated clearance of stress granules may involve interactions with NS1 binding protein (NS1-BP); NS1-BP binds to the UBA domain of p62 [12]. In this context, NS1-BP suppresses ubiquitination of p62 which may stabilize p62 to promote autophagic removal of stress granules [12]. Interestingly, arsenite-induced oxidative stress leads to faster stress granule formation and disassembly in astrocytes than in neurons [94]. These observations raise the possibility that under metabolic stress, p62 may play dual roles in astrocytes, supporting antioxidant defense through the KEAP1-NRF2 pathway and modulating stress granule dynamics.

Our study establishes a framework for understanding how neurons and astrocytes differentially deploy p62 in response to distinct forms of cellular stress. Future investigations will need to define how p62 is balanced between autophagy and the ubiquitin-proteasome system [1, 13]. Primary astrocytes have been reported to exhibit higher proteasomal activity than primary neurons [95], suggesting that these cell types may differ in how proteostatic resources are allocated [33]. Defining how this balance shifts under different stress conditions will be essential for understanding how neurons and astrocytes preserve homeostasis and adapt to metabolic and proteotoxic challenges. More broadly, delineating cell-type-specific p62 stress responses may provide insights into cell type vulnerabilities in neurodegenerative disease.

## CONFLICT OF INTEREST STATEMENT

The authors declare no conflicts of interest.

## ACKNOWLEDGMENTS

This work was supported by NIH grants F31NS132431 to D.K.S., F31NS132453 to M.L.S, and R01NS110716 and Research Supplement to Promote Diversity in Health-Related Research to S.M.. We thank Dr. Mary Putt (Biostatistics and Bioinformatics Core at the Intellectual and Developmental Disabilities Research Center at the Children’s Hospital of Philadelphia) and Dr. Edward Lee (Perelman School of Medicine at the University of Pennsylvania) for advice with statistical analyses. We thank James Shorter and David Kedeme for critical feedback.

## AUTHOR CONTRIBUTIONS

Conceptualization, D.K.S. and S.M.; Methodology, D.K.S. and S.M.; Validation, D.K.S., E.M.S., S.M.; Investigation, D.K.S. E.M.S., M.L.S.; Formal Analysis, D.K.S.; Resources, S.M.; Data Curation, D.K.S., E.M.S., M.L.S., S.M.; Writing – Original Draft, D.K.S. and S.M.; Writing – Review and Editing, D.K.S., E.M.S., M.L.S., M.C.V., S.M.; Visualization, D.K.S., M.C.V., S.M.; Supervision and Project administration, S.M.; Funding Acquisition, D.K.S., M.L.S., S.M..

---

**Figure S1.**
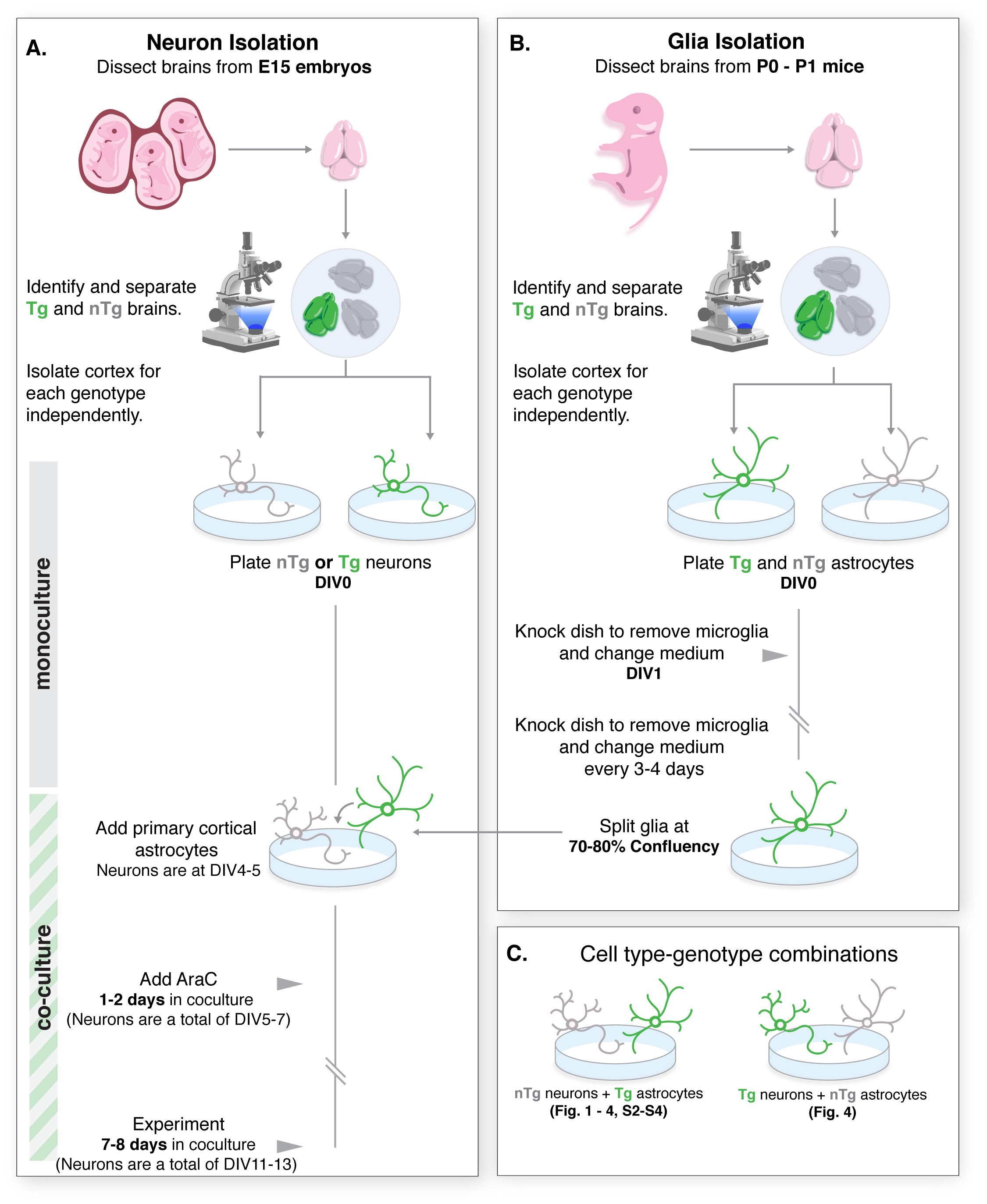
Method for coculturing neurons and astrocytes. **(A)** Cortical mouse neurons (∼96% pure) are isolated from either GFP-LC3 transgenic (Tg) embryos or non-transgenic (nTg) embryos at embryonic day 15 (E15) and plated onto glass coverslips. **(B)** Cortical glia enriched for astrocytes are isolated from P0-P1 pups of the opposing genotype and **(A)** plated onto neurons that are at DIV 4-5. Neurons and astrocytes are then cocultured for 7-8 days (neurons are a total age of DIV11-13 in culture) before performing experiments. **(C)** Cell-type-specific genotype combinations used in this study are denoted within each respective figure.

**Figure S2.**
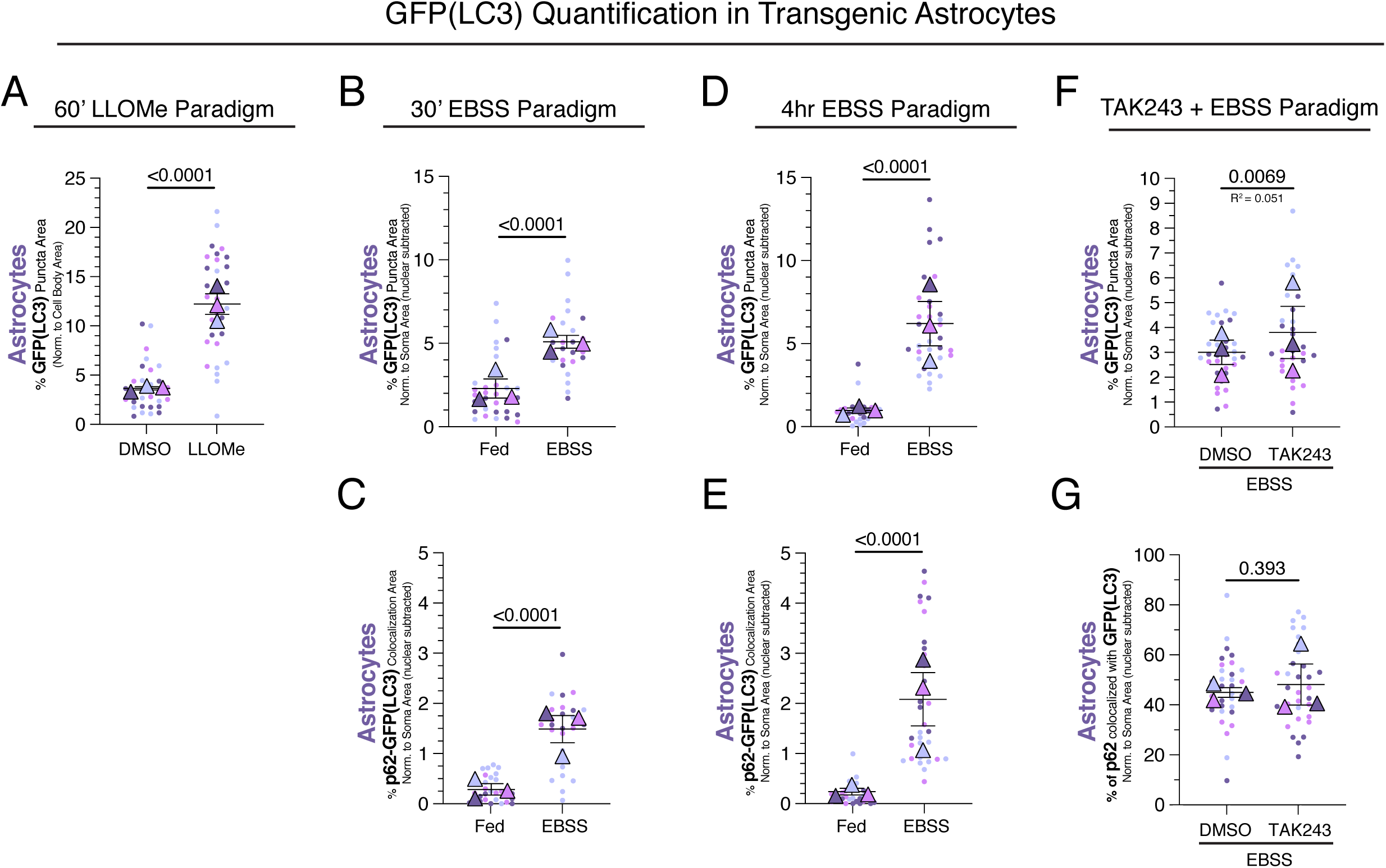
Quantification of autophagosome area in GFP-LC3 transgenic astrocytes under different treatment paradigms. **(A)** Co-cultures of GFP-LC3 transgenic astrocytes and non-transgenic neurons were treated for 1 hr with 1 mM LLOMe to induce lysosomal damage, or an equivalent volume of DMSO solvent as a control. Quantification of total area occupied by GFP(LC3)-positive puncta normalized to soma area in astrocytes. Horizontal bars represent the means of the biological replicates ± SEM; shown is the p-value from a LME model; N=32 astrocytes from 3 independent experiments; 7-8 DIV of coculture. **(B-E)** Cocultures of GFP-LC3 transgenic astrocytes and non-transgenic neurons were treated with EBSS (or regular media as a fed control) for 30 minutes **(B-C)** and 4 hours **(D-E)**. Quantification of area occupied by GFP(LC3)-positive puncta per cell normalized to cytoplasmic soma area of each respective cell **(B, D)**, and colocalization area between p62 and GFP(LC3) puncta normalized to cytoplasmic soma area of each respective cell **(C, E)**. **(B-C)** Horizontal bars represent the means of the biological replicates ± SEM; shown are p-values from a LME model; N= 27-30 astrocytes **(B)** or N= 26-27 astrocytes **(C)** from 3 independent experiments; 7 DIV of coculture. **(D-E)** Horizontal bars represent the means of the biological replicates ± SEM; shown are p-values from a LME model; N= 31-35 astrocytes **(D)** or N= 27-32 astrocytes **(E)** from 3 independent experiments; 7-8 DIV of coculture. **(F-G)** Cocultures of GFP-LC3 transgenic astrocytes with non-transgenic neurons were pretreated for 1.5 hr with 1 µM TAK-243 to inhibit the E1 ubiquitin activating enzyme or an equivalent volume of DMSO solvent as a control, then treated for 30 minutes with EBSS. Quantification of total area occupied by GFP(LC3)-positive puncta normalized to cytoplasmic soma area **(F)** or percentage of p62-positive puncta that colocalize with GFP(LC3)-positive puncta normalized to cytoplasmic soma area **(G)**. Horizontal bars represent the means of the biological replicates ± SEM; shown are the p-values from a LME model; N=32-33 astrocytes **(F-G)** from 3 independent experiments; 7-8 DIV of coculture.

**Figure S3:**
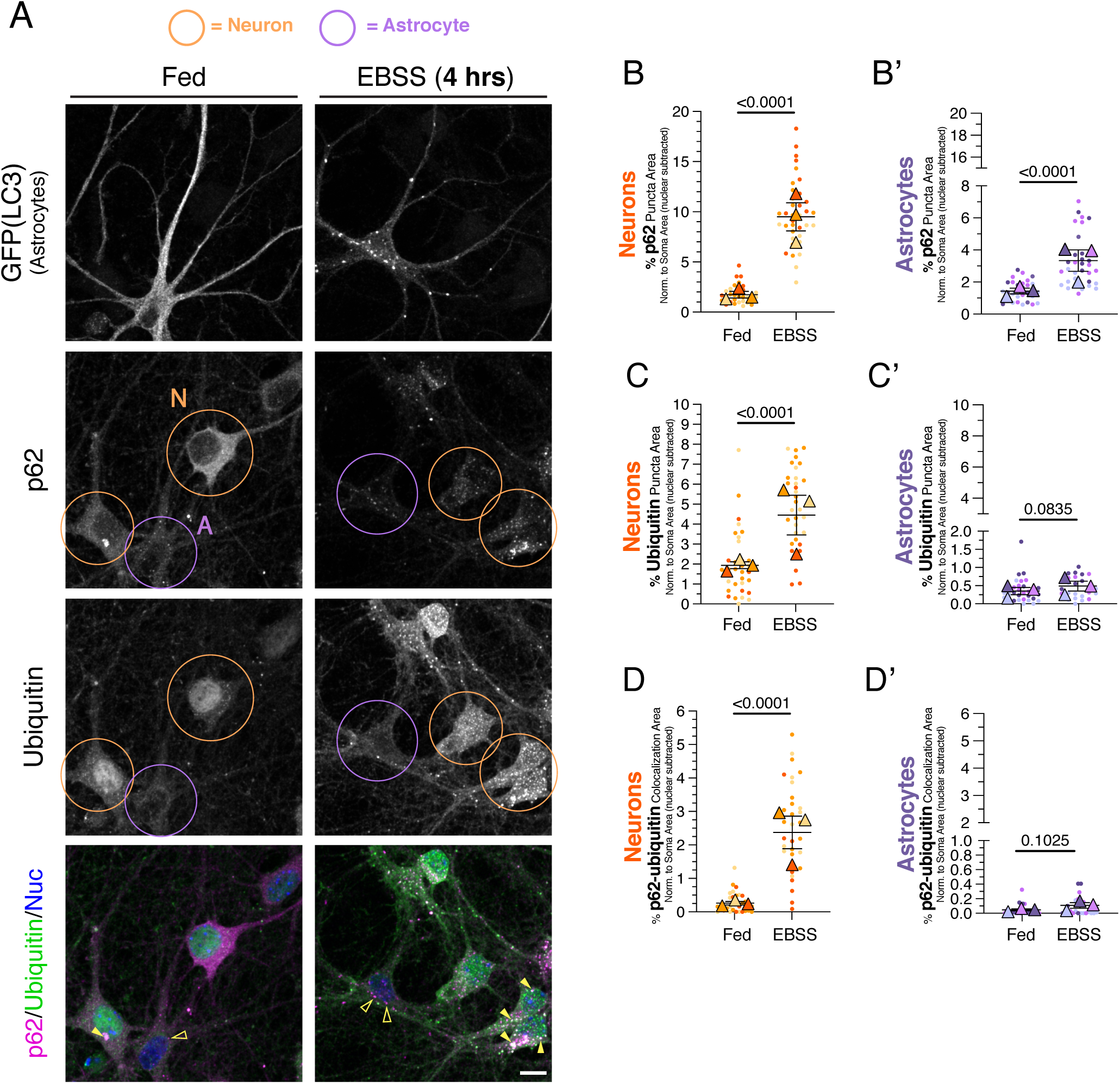
Long-term metabolic stress preferentially increases p62-Ub structures in neurons as compared to astrocytes. **(A)** Cocultures of neurons and astrocytes were treated for 4 hours in EBSS (or regular media as a fed control) and then immunostained for GFP (LC3; labels GFP-LC3 transgenic astrocytes), p62, and ubiquitin; nuclei were labeled with Hoechst. Shown are maximum projections of z-stacks. Images for GFP(LC3), p62, and ubiquitin are grayscale-matched within a protein marker, across treatment conditions. Filled yellow arrowheads indicate colocalization between p62 puncta and ubiquitin puncta. Empty yellow arrowheads indicate p62 puncta with no ubiquitin puncta correlate. Neurons are circled in orange and astrocytes are circled in purple. Scale bar, 10 µm. **(B-B’)** Quantification of total area occupied by p62-positive puncta normalized to cytoplasmic soma area for neurons **(B)** or astrocytes **(B’)**. Horizontal bars represent the means of the biological replicates ± SEM; shown are the p-values from a LME model; N=31-36 neurons and N=30-32 astrocytes from 3 independent experiments; 7-8 DIV of coculture. **(C-C’)** Quantification of area occupied by Ub-positive puncta normalized to cytoplasmic soma area for neurons **(C)** or astrocytes **(C’)**. Horizontal bars represent the means of the biological replicates ± SEM; shown are the p-values from a LME model; N=32-33 neurons and N=24-31 astrocytes from 3 independent experiments; 7-8 DIV of coculture. Images are grayscale-matched within protein marker and across treatment conditions. **(D-D’)** Quantification of total area occupied by p62-positive and ubiquitin-positive puncta normalized to cytoplasmic soma area for neurons **(D)** or astrocytes **(D’)**. Horizontal bars represent the means of the biological replicates ± SEM; shown are the p-values from a LME model; N=30-31 neurons and N=23-26 astrocytes from 3 independent experiments; 7-8 DIV of coculture.

**Figure S4.**
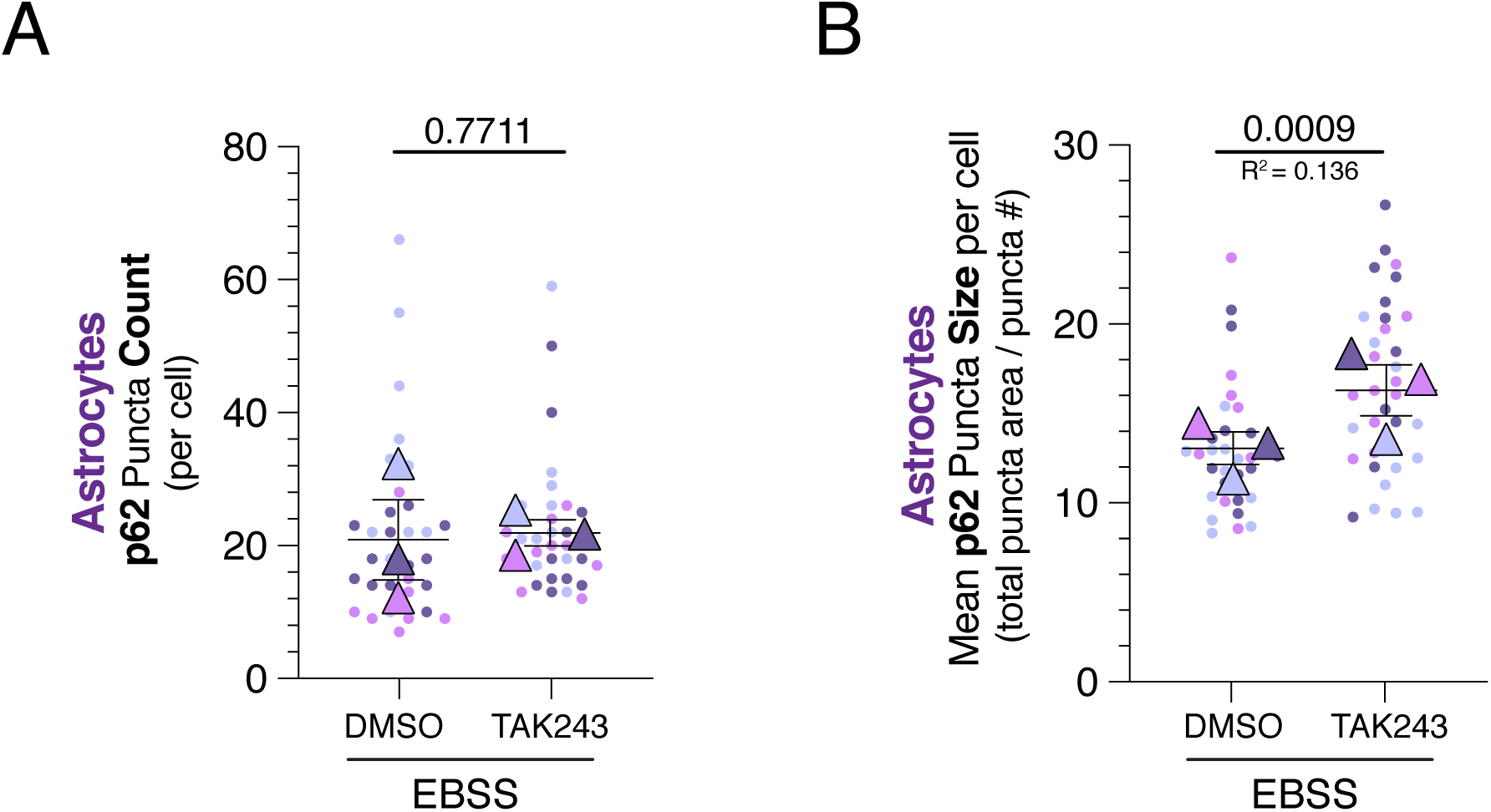
E1 inhibition during metabolic stress only increases p62 puncta size, but not p62 puncta number, in astrocytes. Cocultures of neurons and astrocytes were pretreated for 1.5 hr with 1 µM TAK-243 to inhibit the E1 ubiquitin activating enzyme or an equivalent volume of DMSO solvent as a control. Cells were then incubated for 30 minutes in EBSS (or a fed media control), in the presence of 1 µM TAK-243 or DMSO, to induce nutrient deprivation. Cells were immunostained for GFP (LC3; labels GFP-LC3 transgenic astrocytes), p62, and ubiquitin; shown are maximum projections of z-stacks. Quantification of the number of p62 puncta per cell **(A)** and mean p62 puncta size (i.e., total puncta area divided by puncta number for each cell) **(B)** for astrocytes. Horizontal bars represent the means of the biological replicates ± SEM; shown are the p-values from a LME model; N=32-34 astrocytes **(A-B)** from 3 independent experiments; 7-8 DIV of coculture.

## Notes

### Competing Interest Statement

The authors have declared no competing interest.

